# Ameliorating Emotional Attention through Modulation of Neural Oscillations with Transcranial Alternating Current Stimulation

**DOI:** 10.1101/2021.03.08.434260

**Authors:** Shuang Liu, Yuchen He, Dongyue Guo, Xiaoya Liu, Xinyu Hao, Pengchong Hu, Dong Ming

## Abstract

**Background:** Numerous clinical reports suggest that psychopathy like schizophrenia, anxiety and depressive disorder is accompanied by early attentional abnormalities in emotional information processing. In the past decade, the efficacy of transcranial alternating current stimulation (tACS) in changing emotional functioning has been repeatedly observed and has demonstrated a causal relationship between endogenous oscillations and emotional processing. However, tACS effects on emotional attention have not yet been tested.

**Methods:** A total of 53 healthy participants were randomized to 2 groups, and they were subjected to active or sham tACS at individual alpha frequency (IAF) in the bilaterally dorsolateral prefrontal cortex (dlPFC). Participants and received this treatment for 20 min durations daily for 7 consecutive days. On days 1 and 7, electroencephalogram (EEG) recording of 8 minute resting with eyes open and closed. Responses to a facial emotion identification task were also recorded to measure alpha changes and event-related potential (ERP) alterations.

**Results:** On day 7 after tACS, the active group showed a more clear elevation in alpha power at rest, especially in open state around stimulation area, compared to the sham group. ERPs revealed a significant larger P200 amplitude after active stimulation (*p* < 0.05), indicating attentional improvement in facial emotion processing. Additionally, a notable positive correlation (*p* < 0.05) between alpha power and P200 amplitude was found, providing an electrophysiological interpretation regarding the role of tACS in emotional attention modulation. In addition, the IAF-tACS showed an obvious advantage in alpha entrainment compared to an additional 10 Hz-tACS.

**Conclusions:** These results support a seminal outcome for the effect of IAF-tACS on emotional attention modulation, demonstrating a feasible and individual-specific therapy for neuropsychiatric disorders related to emotion processing, especially regarding oscillatory disturbances.

## Introduction

Emotion plays an important role in daily life, and emotion processing is a crucial skill for adaptive social behavior. Recently, emotion processing has been under intensive focus in the field of human neuroscience (Camacho, Karim, & Perlman, 2019; Dolcos et al., 2020). Clinical research indicates that the phenomenon of numbness to the emotional information for psychopathy like schizophrenia, anxiety and major depressive disorder may be explained by early attentional abnormalities sometimes (Connell, Danzo, Magee, & Uhlman, 2019; Martínez et al., 2019; Schulze, Schulze, Renneberg, Schmahl, & Niedtfeld, 2019; Wieckowski, Capriola-Hall, Elias, Ollendick, & White, 2019). For instance, patients with autism spectrum disorders tend to exhibit deficits in early stages of facial emotions processing at the neural level, which undermines their subsequent processes and behaviors during emotion-related tasks (Bunford, Kujawa, Fitzgerald, Monk, & Phan, 2018; Sokhadze et al., 2015). As the initial stages of emotion processing, early attention to emotional cues always occurs prior to emotional information extraction and contributes to the subsequent processing (i.e., emotional recognition, evaluation, and cognitive control processing). However, the mechanisms underlying early attentional stage for emotion processing remain unclear, which necessitates preclinical research with healthy subjects to access effective treatments and provide further insights into clinical applications.

As one of the classical approaches used in emotion processing research, facial processing marked by rapid attentional processing has caused great concern due to distinct responsiveness of early components to faces (Jessen & Grossmann, 2020; Rischer et al., 2020). Due to its excellent temporal resolution and sensibility to specific psychological processes, the time course of facial emotion processing has been investigated mostly using event-related potentials (ERPs) and witnessed an evident rise to inspect neurophysiological processes associated with emotional processing in the last two decades. ERPs can reveal underlying components elicited by emotional stimuli that are analyzed in posterior (parietal) areas and reveal salient alterations in amplitude or latency. This reflects the degree to which component-related psychological process are engaged and the time point of completion for the psychological operation respectively (Campanella et al., 2002). In particular, early ERP components (100–300 ms), such as N100 and P200 usually relate to attentional processes and are considered as indices of early attention to emotional stimuli. In contrast, later components (>300 ms) such as P300 have been thought to reflect more elaborative, cognitive and perceptive, top–down, emotional information processing, including face feature encoding and emotional evaluation (Fields & Kuperberg, 2012). Therefore, exploring the potential relationship between early ERP components in response to facial stimuli and psychological processes during initial attentional stages in response to emotional tasks is of great importance.

Apart from dynamic course measurements using ERPs, analysis in frequency domain have also contributed to uncovering brain mechanisms related to the cerebral cortex and emotion processing (Bocchio, Nabavi, & Capogna, 2017; Likhtik & Johansen, 2019). Previous electrophysiological studies have demonstrated associations by correlating emotion processing and brain oscillations. In particular, alterations or disturbances in particular oscillations can be linked to psychiatric disorders (Kang et al., 2018; Sohal, 2012). In some cases, brain oscillations are amenable to external intervention and can be experimentally manipulated though neuromodulation techniques, which has gained popularity when combined with electroencephalography (EEG) in cortical rhythm research (Yavari, Jamil, Samani, Vidor, & Nitsche, 2018). Transcranial alternating current stimulation (tACS) is noninvasive stimulation paradigm that has potential to entrain spike timing at single-unit level and facilitate endogenous cortical oscillations via employing weak sinusoidal electric current to the scalp (Johnson et al., 2020; Krause, Vieira, Csorba, Pilly, & Pack, 2019; Liu et al., 2018; Polania, Nitsche, & Ruff, 2018). Much of the recent interest in tACS stems from its specific frequency manner, which offers the possibility to entrain targeting oscillation and thus characterize different cognitive processing activity in specific brain regions (Hanslmayr, Axmacher, & Inman, 2019; Krause et al., 2019; Lafon et al., 2017). Abundant studies have suggested its efficacy in changing brain function and active use in the treatment of psychiatric diseases such as MDD and anxiety. Thereby, it has become a strong candidate for clinical adjuvant therapy over the past several decades (Alexander et al., 2019; Clancy et al., 2018; Shekelle et al., 2018; Singh et al., 2016). However, little is known about the effects of tACS on early attentional stage for emotion processing. It is an emerging issue whether tACS can modulate emotional attention by entraining endogenous neural oscillations.

Two factors are key in addressing the hypothesis above: frequency band and brain regions. Alpha band (8–13 Hz), known initially as “idling rhythm,” operates through power alternation and sustains higher intrinsic cortical activity (Zheng et al., 2019). Vast empirical evidence links alpha oscillations with the activity of emotional and attentional systems during tasks associated with affective processing as well as rest state studies (de Aguiar Neto & Rosa, 2019; Eidelman-Rothman, Levy, & Feldman, 2016; ter Huurne et al., 2013). In particular, aberrant alpha behavior has been investigated by numerous clinical studies. Depression-related alpha abnormalities are mostly centered at frontal and posterior alpha asymmetry at-rest (Jesulola, Sharpley, Bitsika, Agnew, & Wilson, 2015). Patients with bipolar disorder have shown inconsistent alpha power alternations compared to healthy participants, and this asymmetry varies among different tasks (Harmon-Jones et al., 2008). Studies assessing alpha power in anxiety, and post-traumatic stress disorders underline the relationship between abnormalities in alpha activity and psychopathology (Crost, Pauls, & Wacker, 2008; Imperatori et al., 2014). In general, studies assessing alpha activity often focus on fixed or individualized alpha bands, and the latter has been suggested to be more advantageous, since it takes into account an individual’s alpha characteristics (Bazanova & Vernon, 2014). Considering together, these observations indicate the value of alpha functionality in attentional and affective processing.

As for the target brain region, there is robust evidence to indicate that the prefrontal cortex is involved in emotion-related functions. As part of the dorsal system, the dorsolateral prefrontal cortex (dlPFC) plays a central role in cognitive regulation and executive function of emotion, including effortful regulation of affective states, which is key in processing stimuli without self-referential emotional content (e.g., facial expression and visual scenes) (Dixon, Thiruchselvam, Todd, & Christoff, 2017; Raschle et al., 2019; Zhang, Japee, Safiullah, Mlynaryk, & Ungerleider, 2016). Similar to alpha activity, human neuroimaging and electrophysiological studies have provided great insight into the link between dlPFC abnormalities and psychiatric disorders (Liston et al., 2014). Although explicit correlation between dlPFC deficit and emotional function has been well-established, it remains partially unclear how the dlPFC activity affects emotion processing at the neural level.

Although tACS of specific frequency bands have recently been investigated in various domains, few studies have considered alpha-tACS in the dlPFC as an intermediate to explore their interactive connections, which may represent underlying physiological mechanism in emotion processing. Given the emotional function of the dlPFC and attention-affective role of alpha rhythm, this has suggested the notion that alpha-tACS in the dlPFC is a powerful tool to promote emotional behavior. It also promotes the investigation of early attentional stages during emotion processing, which is the pioneer preclinical research in the field.

Described here is a single-blind and sham-controlled experiment in healthy subjects to validate tACS-effects of this promising emotional attention aspect, which contains two separate tasks of at-rest and emotion identification. tACS was delivered over the bilateral dlPFC based on individual alpha frequency (IAF). It was hypothesized that the effects of tACS on emotional attention will provide a potential regulation approach regarding emotion processing.

## Methods and Materials

### Participants

A total of 53 right-handed healthy participants (27 females, mean age = 23 years, age range = 20 to 27 years) participated in the study. All of the participants, with normal or corrected to normal vision, gave written informed consent and reported no history of neurological or psychiatric disorders or prior transcranial electrical stimulation experience. Prior to the experiment, participants were instructed to complete mood questionnaires using the State-Trait Anxiety Inventory (STAI; Spielberger et al., 1968) and Difficulties in Emotion Regulation Scale (DERS; Gratz & Roemer, 2004) to access self-reported emotion regulation functioning (Bardeen, Fergus, & Orcutt, 2012; Barnes, Harp, & Jung, 2002).

A total of 53 participants were randomized to one of two groups: active group (26 participants) or sham group (27 participants). Both groups showed stable normal levels on the questionnaires and revealed no salient difference in questionnaire and age. EEG data of two subjects from the active group were omitted because of unusually large differences at baseline due to instability during recording.

### Procedure

The experiment involved a randomized, single-blind, sham-controlled design and was performed in an electrically shielded and sound-attenuated cabin. All of the procedures were conducted with approval from the Institutional Review Board at Tianjin University and ethics committee of Tianjin University Tianjin Hospital.

The procedure was clearly explained to all participants beforehand. The entire procedure required 7 days (Figure 1A). EEG recording was executed repeatedly on day 1 (before stimulation) and day 7 (after stimulation). During EEG recording, each participant underwent two tasks, including 8 minute of resting EEG with eyes open and close task and facial emotion identification task. In each task, the participants were asked to relax, minimize their body movement to reduce the appearance of relevant artifacts in the EEG recordings, and concentrate.

**Figure 1:**
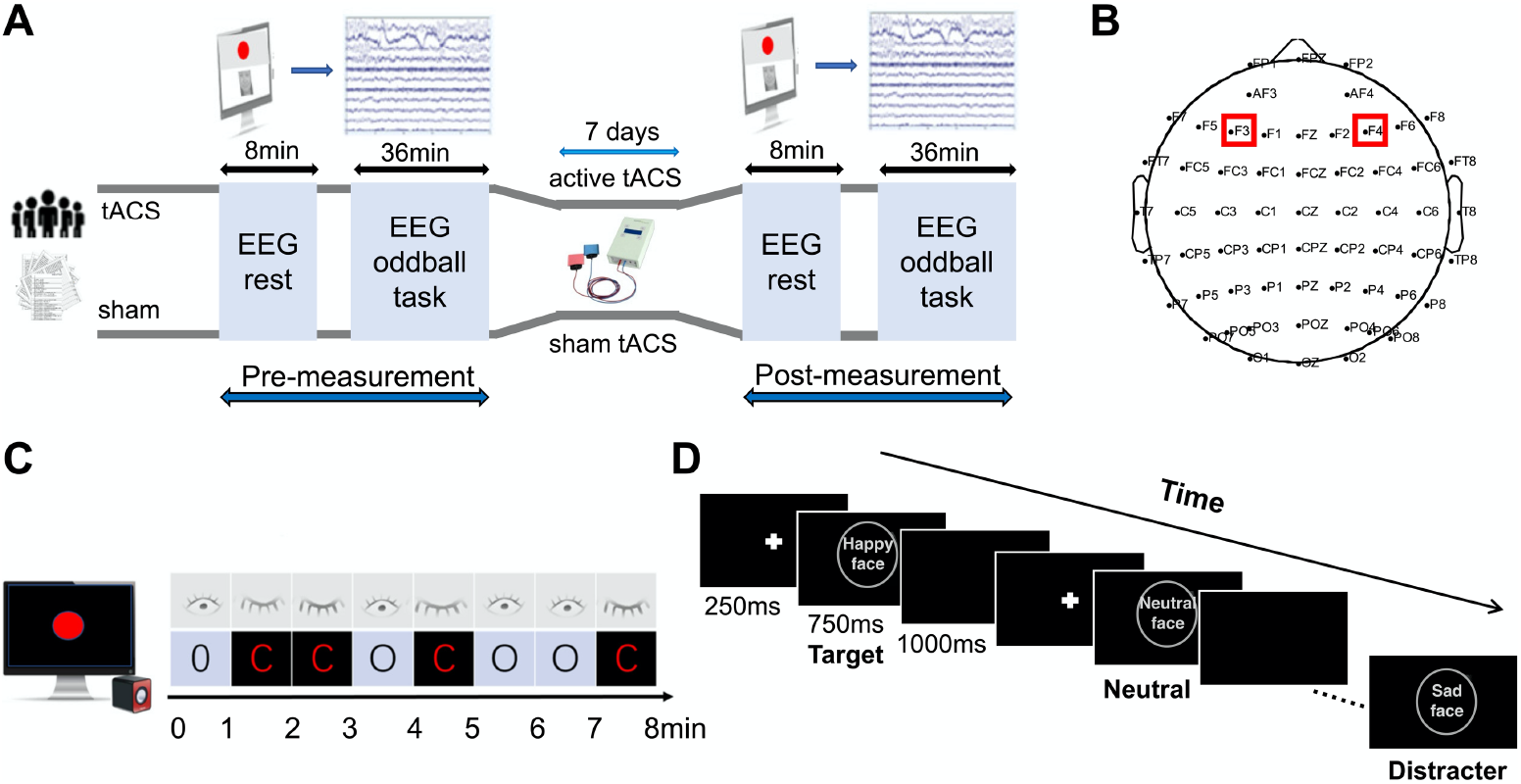
Experimental paradigm. (A) Experimental procedure. (B) EEG and tACS electrode placement on the scalp. (C) Illustration of 8 minute resting task with eyes closed (C) and open (O). The periods consisted of eight 1-minute intervals of rest in the following order: OCCOCOOC. Participants concentrated on the red circle on the screen in the open intervals. After the last interval, participants heard “beep” sound when the red circle turned blue, indicating the end of the procedure. (D) Illustration of facial emotion identification task. Shown is a possible order of pictures in the positive condition, with each stimulus presented in the center of the screen for 1 second, including 250 ms for fixation cross and 750 ms for face stimulus, followed by a blank screen for 1000 ms. The task was preceded by a short practice period. In each block, participants kept their eyes fixated on the screen and responded as quickly as probable to the target stimuli by pressing the space bar and ignoring distracter stimuli as well frequently presented standard stimuli. To avoid the leakage of identifying information of people, the actual facial stimuli were replaced with graphics in the flowchart.

### tACS

tACS was administered through a battery-operated stimulator (DC Stimulator, Neuroconn, Germany) and applied in sinus pattern with the initial phase of 0, which is deemed as the optimal phase difference for task performance (Kanai, Chaieb, Antal, Walsh, & Paulus, 2008). Two round rubber electrodes (2.5 cm diameter) in saline-soaked sponges were placed bilaterally over F3/F4 of the international 10–20 system (Figure 1B). The impedance was kept below 10 KΩ of stimulation duration. Stimulation intensity was 1.5 mA (peak-to-peak), below individual phosphene and discomfort threshold. Stimulation frequency was individual alpha frequency (IAF, see Supplement) determined for each participant on day 1, given that alpha frequency varies among age, memory and genetic background. Each participant received tACS for 7 consecutive days for a duration of 20 minutes each day at about the same time (± 2 hour). Regardless of the authenticity of the stimulation, all participants were required to remain relaxed during the stimulation process.

### Tasks

For the resting task, EEG acquisition started with 8 minute of resting EEG. During recording, participants were instructed to remain still and relaxed with their eyes open (O) and closed (C) in two alternating orders following the voice prompts (Figure 1C), with the goal of averaging the difference between open and closed signals (Adolph & Margraf, 2017). For the facial emotion identification task, a well-established visual oddball paradigm was used to access neurophysiologic effects of incidental emotion processing and retrieve ERP components (Figure 1D). Participants were instructed to complete six blocks made of three positive and three negative conditions. The block order was random for each participant. Each block lasted for 6 minutes and consisted of a pseudorandom series of 180 pictures with positive, negative and neutral faces from the Chinese Affective Picture System (CFAPS). Within each block, facial stimuli were presented with overall proportions of 70% standard stimuli, 15% targets and 15% distracters. Facial stimuli with emotional valence opposite to the targets were used as distracters (e.g., positive distracters in the block with negative targets), and neutral faces were used as standard stimuli.

### EEG

Raw EEG data was acquired by a SynAmps2 amplifier using a 64-channel EEG acquisition system (NeuroScan, Germany) sampled at 1,000 Hz. The right aural tip was served as reference and AFz was served as ground. The analysis included preprocessing, power spectral estimation, and ERP analysis. For spectral analysis, the resting EEGs were separated according to the eyes-open and eyes-closed state, then epoched into 5 second segments, after which power spectrum was estimated with a 1024-point fast Fourier transform (FFT) using Welch’s method (0.5 Hz resolution). The resulting spectrum was subsequently averaged across epochs as well as across the individual alpha band. To determine spectrum changes between day 1 and day 7, absolute power (μV^2^/Hz) in the fixed alpha frequency band (8–13 Hz) and individual alpha band (IAF ± 2.5 Hz) were calculated. Notably, the absolute power of alpha band is the integral of all power values within the frequency range. However, normalized spectrum at electrode level were accessed for each subject, with the aim to eliminate individual differences within each group and compare obvious changes in alpha power across groups.

Then, ERP elicited by positive, negative, and neutral faces were analyzed using the signals recorded during facial emotion identification task. In this study, N100 (60–120 ms), P200 (120–200 ms) and P300 (350–550 ms) components were measured by peak value (baseline to peak of each component) and latencies (from stimulus onset to the peak) at Pz electrode, based on grand average ERP activity and previous studies. The specific method for preprocessing and ERP analysis can be found in the Supplement.

## Results

### Alpha spectrum

To explore the effects of tACS on neuronal entrainment, changes in resting-state power spectrum in individual frequency band of all-channel resting EEG were assessed. The deviation in power spectrum was defined as relative changes calculated by the post- minus the pre-stimulation normalized data. Compared to 8 minute resting data, relative power spectrum changes in the individual alpha band were obviously enhanced in open and close states after stimulation and were concentrated around IAF for active group, while only enhanced in the open state for the sham group (Figure 2A, 2B). Besides, there were no significant IAF differences found between the two groups.

**Figure 2:**
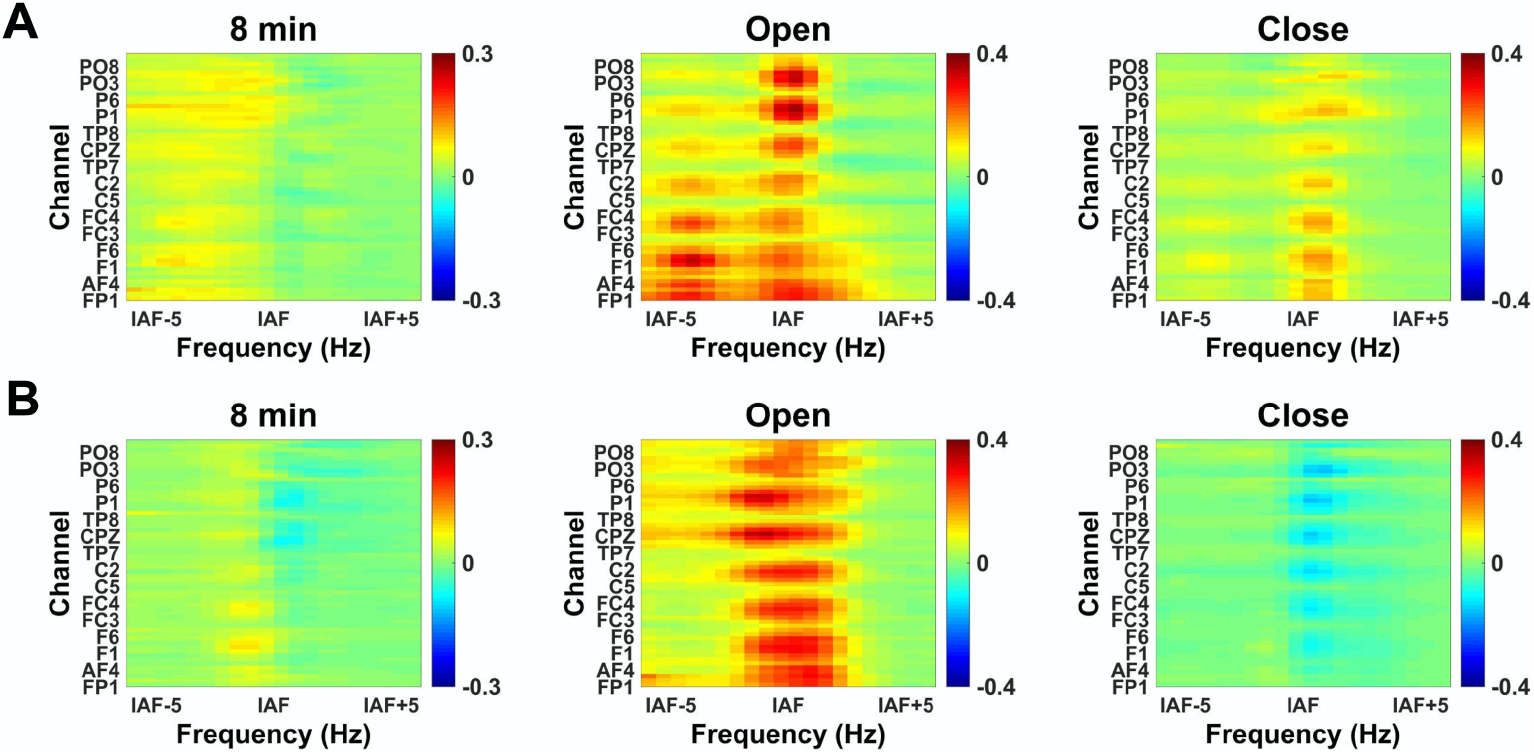
Relative changes in mean normalized alpha spectrum of full 8 minutes resting, open, and close states across subjects within channels in individual alpha bands for (A) active and (B) sham groups. The left column shows spectra of 8 minute resting EEG, the middle column shows spectra of the open state, and the right column shows the spectra of the close state. The x-axis shows individual alpha band (IAF ± 5Hz). To clearly display channel distribution, the y-axis shows only 15 channels.

To further verify alpha changes after stimulation in open and close states, alpha power deviation of normalized spectrum in fixed and individual alpha band were subsequently computed. Paired t-tests using pre- and post-stimulation data at the electrode level were then performed, and the resulting topographies for the two groups are illustrated in Figure 3A. In the active group, the comparison to pre-stimulation spectra indicated that alpha activity was clearly modulated in the open state and exhibited a significant elevation in the prefrontal cortex (*p* < 0.05), while there was only a slight enhancement in the close state. Regarding the sham group, the alpha spectrum was elevated from the anterior to posterior direction in the open condition and differed subtly in the close state, with a significant change in the occipital lobe. To account for the results in the sham group, the power of each individual was analyzed, and it was found that 12 of 27 participants in the open state and 10 in the close state exhibited increased alpha power after tACS, revealing a random trend compared to the active group.

**Figure 3:**
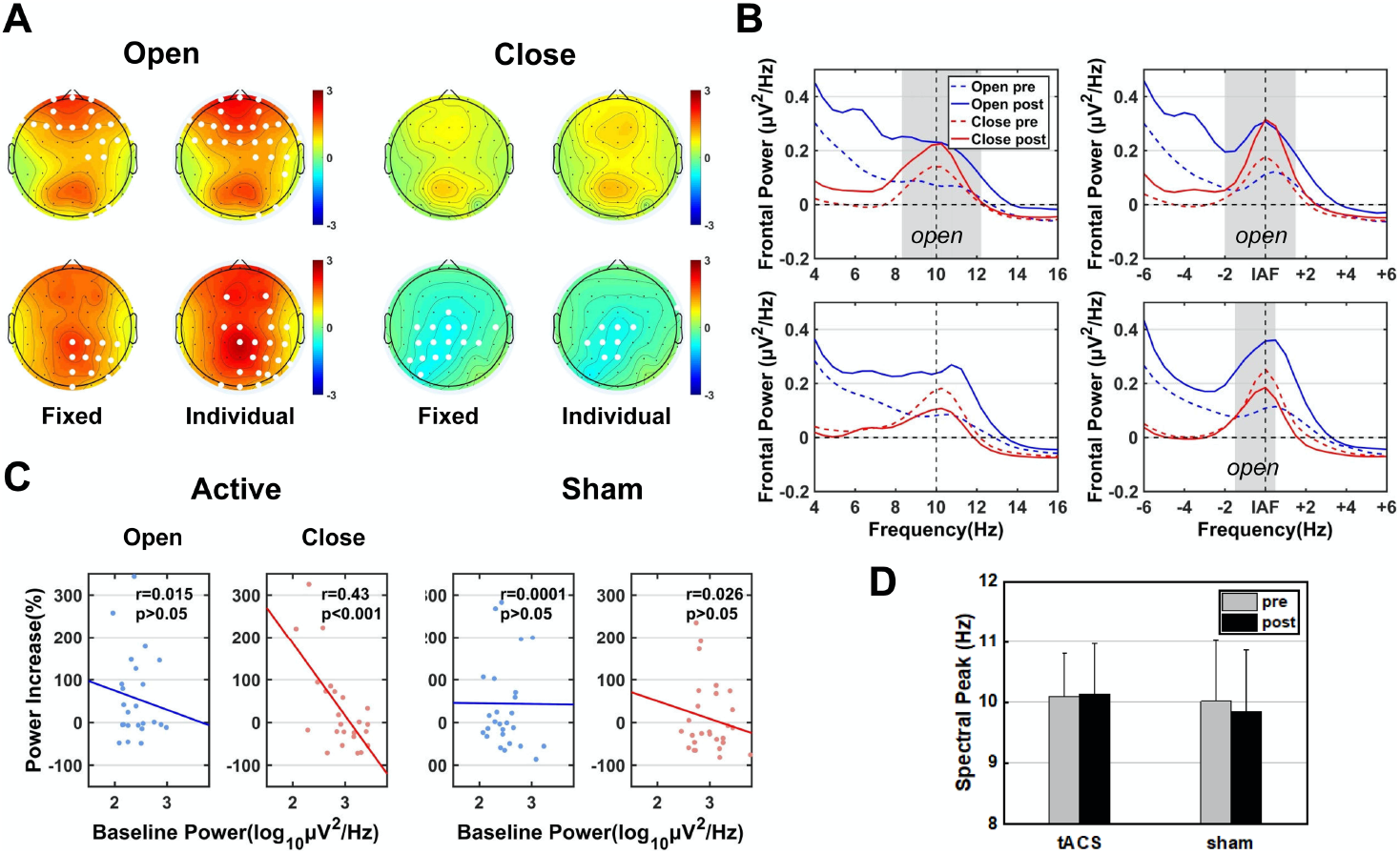
Alpha activity changes after 7 day tACS in open and close states. (A) Scalp topography of alpha power changes in active (top) and sham (bottom) groups. The left column shows power changes in fixed alpha band (8–13Hz) and the right column shows power changes in individual alpha band (IAF ± 2.5Hz). White dots in the topography map indicate significant channels, *p* < 0.05. (B) Grand mean normalized frontal power across subjects in fixed (left) and individual (right) bands. Frontal power is the integral of all power values within the frontal electrodes. The top row shows frontal power in the active group and the bottom row shows the sham group. Open pre = frontal power of open state before stimulation; Open post = frontal power of open state after stimulation; Close pre = frontal power of close state before stimulation; Close post = frontal power of close state after stimulation. Gray shade represents significant frequency interval in the open state (*p* < 0.05). (C) Linear correlation analyses of the absolute alpha power on day 1 and the individual alpha power increases in open (blue) and close (red) states. Each dot depicts one subject. Significance of Pearson correlation (p) and R-square (r) are marked in upper left corners of the scatterplots. (D) Spectral peak analyses in alpha band of two groups. Pre = peak frequency in alpha band of resting EEG before stimulation; post = peak frequency in alpha band of resting EEG after stimulation.

Considering the diffusion of alternating current, frontal alpha changes in the open and close states were analyzed (Figure 3B). The frontal alpha power in the active group was clearly elevated in two states and showed a broad significant frequency band in open states (paired t-test, *p* < 0.05), while there was a narrow significant frequency band of individual alpha power in the sham group (*p* < 0.05), which may have been caused by a placebo effect. In accordance with topographies of alpha power, this finding also confirmed tACS had an effect on the open state. Besides, both groups revealed a more concentrated distribution and higher alpha power in individual alpha band centered on the IAF frequency of each participant.

In addition, a significant negative correlation in the close state was found between the log-transformed absolute alpha power on day 1 and the relative increase in alpha power after stimulation (r = 0.43, *p* < 0.001, Figure 3C). This was not observed in the open state (r = 0.015, *p* > 0.05) or sham group (r = 0.026, *p* > 0.05). This indicated that tACS-induced power enhancement was limited by initial alpha power in the close state. Figure 3D shows the tACS-effect of IAF. The peak frequency of the alpha band in the active group was unshifted after stimulation compared to the sham group, suggesting a “locking effect” of neural activity in the stimulation band.

### ERPs

ERP waveforms of target, distracter and neutral stimuli were elicited and illustrated into four parts (two groups, two conditions, Figure 4), with time window expression for each component. As depicted, the oddball paradigm exhibited typical visual ERP waveforms and retrieved three highly predictable ERP components of N100, P200 and P300. It should be noted that the P300 component was induced only by the target and distraction stimuli. To assess tACS effects on ERP modulation, we computed peak amplitudes and latency differences were computed for each component (i.e., spectrum deviation computation), and they are depicted in a histogram. Paired t-tests using data from pre- and post-stimulation were performed. First come to the amplitude parameter, the N100 component showed an almost identical decrease in the two groups and significant for most neutral and distracter stimuli in both positive (neutral: *p* < 0.005, distracter: *p* < 0.01 for active; neutral: *p* < 0.01, distracter: *p* > 0.05 for sham) and negative conditions (neutral: *p* < 0.005, distracter: *p* < 0.01 for active; neutral: *p* < 0.01, distracter: *p* < 0.05 for sham). One possible explanation to account for this finding was increased familiarity of the task after the first experiment.

**Figure 4:**
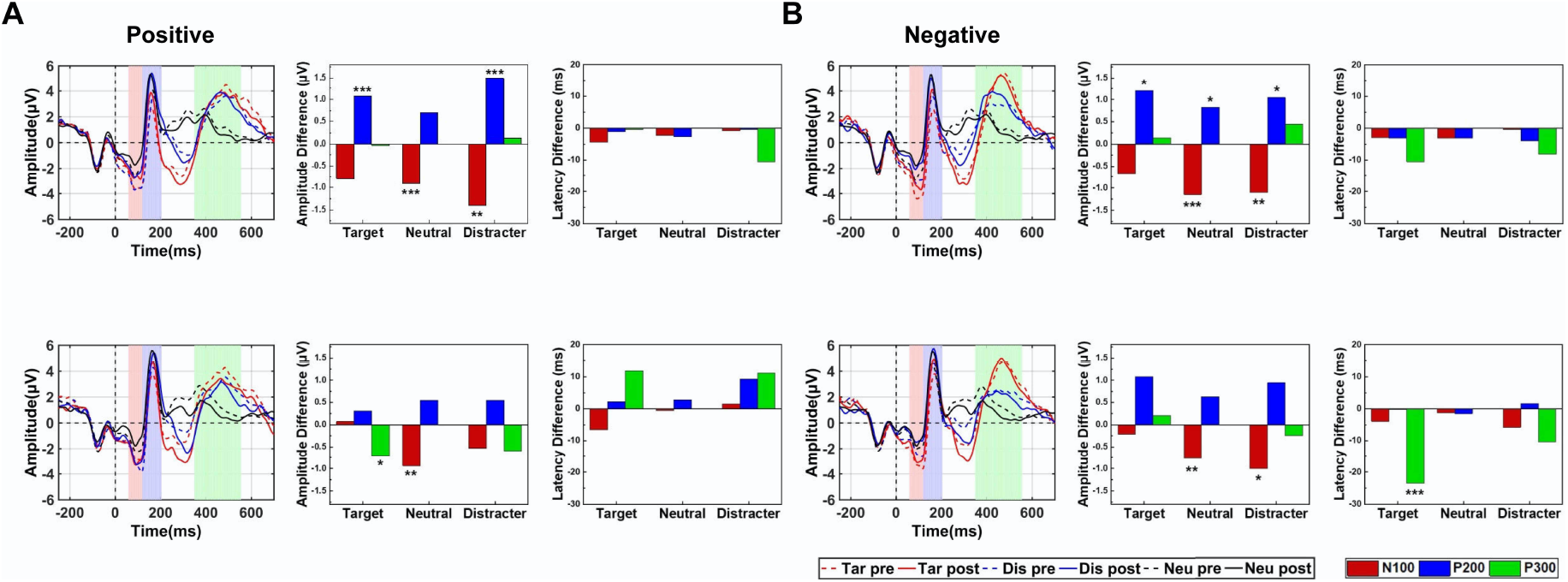
ERPs generated by target, distracter, and neutral stimuli of the two groups under (A) positive and (B) negative condition. Images in the top row show ERP measurements in the active group, and the bottom row shows the ERP measurements in the sham group. (A) Shown is an example of a positive condition in the active group. The left column shows the ERP waveform of three types stimuli, the middle column shows mean amplitudes differences in N100, P200, and P300 components, and the right column shows mean latency differences in the three components. The shaded area in waveforms represent the time windows for each component. Tar pre = waveform elicited by target stimuli before stimulation; Tar post = waveform elicited by target stimuli after stimulation; Dis pre = waveform elicited by distracter stimuli before stimulation; Dis post = waveform elicited by distracter stimuli after stimulation; Neu pre = waveform elicited by neutral stimuli before stimulation; Neu post = waveform elicited by neutral stimuli after stimulation. (* represents the significant different, \**p* < 0.05; \*\**p* < 0.01; \*\*\**p* < 0.005).

The P200 component was significantly elevated for active group in both positive (target: *p* < 0.005, distracter: *p* < 0.005, neutral: *p* > 0.05) and negative conditions (target: *p* < 0.05, distracter: *p* < 0.05, neutral: *p* < 0.05) but was not substantial in the sham group. This indicates that participants in the active group showed strong promotion of early attention of facial emotions after stimulation compared to the sham group. For late ERP component P300, there were slight differences after stimulation between the two groups, in contrast with the above components. These findings suggest that the ERP amplitude modulation of tACS was observed in early components rather than late course processes.

Regarding latency, there was a small decrease in both conditions for the active group, significant results were not observed in the sham group as well. Taken together, the results suggest the effective role of tACS on emotional attention, which was mainly embodied in P200 amplitude modulation rather than latency.

### Correlation analysis

To explore a potential correlation between ERPs and at-rest neural oscillations, both correlation and regression analyses were conducted via Pearson correlations, which is known as a pervasive sample correlation coefficient (r). Considering the main outcome of ERPs, P200 peak value, as well individual alpha power pre- and post-tACS, from the two groups were integrated to create the scatter plot and measure linear relationships. More specifically, correlations based on absolute individual alpha power of frontal cortex and mean P200 amplitude of different stimuli were calculated (Figure 5).

**Figure 5:**
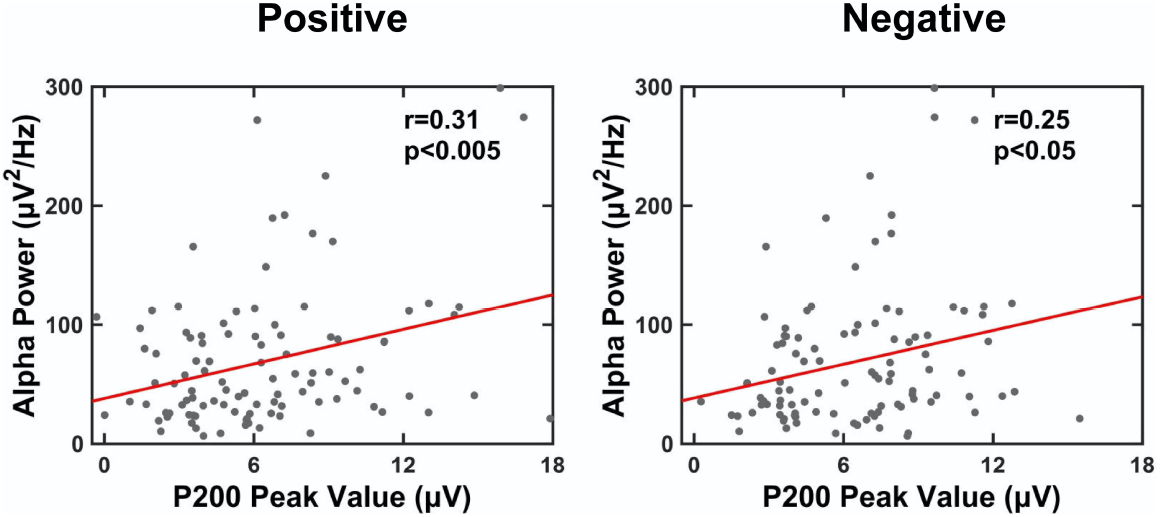
Linear correlation analysis between absolute individual alpha power (y-axis) and P200 peak value (x-axis) in positive (left) and negative (right) conditions. Each scatterplot represents one subject value from two sets of data. Regression lines are drawn in red lines. Significance of Pearson correlation (p) and R-square (r) are marked in the upper left corners of the scatterplots.

A considerably significant and positive correlation was found between P200 peak value and alpha power both in positive and negative conditions. Concerning the enhanced alpha after active tACS mentioned above, the analyses indicate that the higher P200 peak value was largely associated with promoted alpha oscillation, which provides a possible electrophysiology interpretation for the role of tACS role in emotional attention modulation.

## Discussion

Overall, compared to the sham group, participants in the active group displayed increased alpha oscillatory patterns in the frontal lobe at rest, especially in the open state. ERP results revealed a unique increased P200 amplitude in the active group after stimulation. Another finding was a significant positive correlation between frontal alpha power and P200 peak value. These results not only support the neuromodulatory role of the dlPFC in the emotional attention processing but also provide strong evidence that tACS (based on individual alpha activity) can improve the capacity of emotion recognition at early stage by modulating alpha oscillations, which helps underpin the clinical potency of tACS in neuropsychiatric disorders related to emotion processing.

Based on previous evidence from ERP studies, an hypothesis suggesting that there are three stages of facial expression processing has been established (Luo, Feng, He, Wang, & Luo, 2010). In the first stage, perceptual priming processing occurs with temporal characteristics of the N100 component. In this phase, familiarity can facilitate the automatic processing of facial information, which accounted for the significant decline in N100 amplitude in the two groups in this study (Bornstein, Arterberry, & Mash, 2013; Goto & Tobimatsu, 2005). Afterward, emotional information processing starts early attention to emotional stimuli within around 200 ms after stimuli onset. Distinct P200 has supported the association with initial attentional control, which has become a biomarker of attentional impairment for patients with depression (Hu et al., 2017).

An emotional vocalizations research suggested that priming effects were tagged by P200 for both familiar and unfamiliar voices, which excludes the influence of familiarity on P200 amplitude (Föcker, Hölig, Best, & Röder, 2011). With these results, a significant elevation in P200 amplitude in response to distracted stimuli was found in both conditions, indicating that the improved response of emotion recognition for facial emotion had nothing to do with emotion types. For the third stage, there is emotional differentiation among various facial expressions and further evaluation related to corresponding affective valence. Based on the above findings, the results report remarkable effects of tACS on emotional attention modulation, instead of automatic or complex affective processing in the late period.

To our knowledge, this is the first time tACS has been used for studying early attention modulation of emotion processing, demonstrating that tACS may be a potential and feasible treatment in psychiatric population. The study also provides evidence that neuronal oscillations involved in specific neuropsychiatric disorders are associated with disturbances in other brain oscillatory, versus sensory stimulation. Compared to animal models, these physiological effects in healthy humans have greater possibilities for clinical translation because of the specificity of human neocortex, where endogenous ongoing activity in complex oscillations might either amplify or suppress the effects of alternating fields externally (Johnson et al., 2020).

Regarding resting performance, alpha alterations in the open state were more sensitive to tACS intervention, at which stimulation frequency originated. Although both groups revealed an enhanced alpha activity after stimulation, the blatant elevation in alpha power in the active group was focused in the stimulation area. This differed from the posterior cortical areas, where there was higher alpha power demonstrated in the sham group. Interestingly, alpha activity was enhanced in participants with lower endogenous alpha activity, and subtle or even adverse alternations were found in participants with high endogenous alpha activity after stimulation. However, under conditions of higher endogenous alpha power, tACS failed to substantially affect alpha power in the close state and exhibited small increases after tACS (only in the active group). In line with a previous study, greater intensity is required to elevate alpha oscillations when a participant’s eyes are closed (Neuling, Rach, & Herrmann, 2013).

In light of individual differences, tACS was administered based on individual alpha activity. As expected, results revealed a more obvious elevation in individuals compared to the fixed alpha band. This finding is consistent with recent research on tACS, emphasizing its capability to entrain the activity of individual neurons in the primate brain (spatial- and frequency-specifically) (Krause et al., 2019). To verify the effectiveness of tACS based on individual activities, tACS was administered at 10 Hz to 12 additional participants from the same population, as described above. Then, alpha performance in the open state was tested, given the origin of IAF detection (Supplement), and alpha alternations in the open state of the two active groups were compared (Figure 6). In both fixed and individual frequency bands, there was sharp alpha entrainment in the IAF group compared to 10 Hz tACS group under the same stimulation intensity. This phenomenon may be explained stimulation frequency selection of IAF, where corresponded to native regularity pattern of individual network. This finding provides a direct and robust evidence for parameter optimization of tACS.

**Figure 6:**
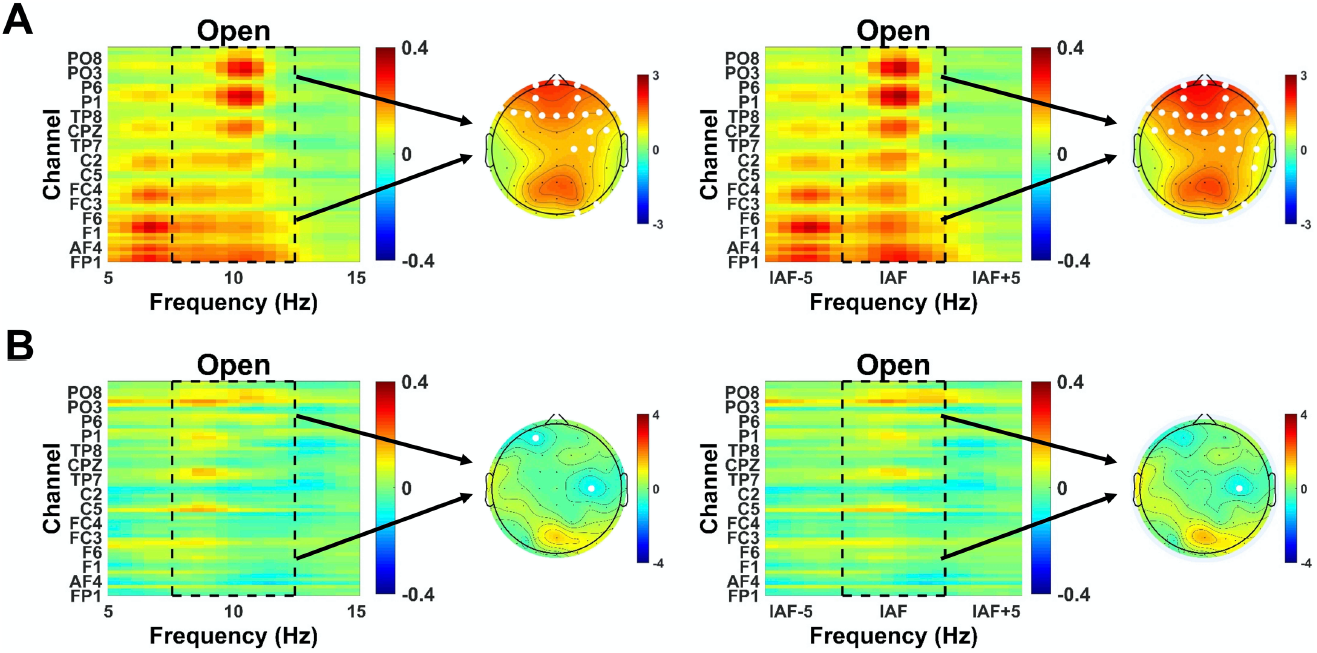
Relative changes of mean normalized broad spectrum and alpha power changes in the open state of IAF-tACS (top) and 10 Hz tACS group (bottom). The left column shows spectrum and alpha power changes in fixed band and the right column shows changes in individual alpha band. The dashed box is the frequency range for alpha power calculation. White dots in the topography map indicate significant channels (*p* < 0.05).

Taking the positive correlation between resting alpha power and P200 value into account, it was found that the improved emotional attention may be explained by alpha entrainment via tACS. This finding relates individual brain oscillation at rest to brain activity during a task-related response. Taken together with the clinical avenue of P200 above, these novel findings verify the important role of resting EEG patterns in emotional research, which has also been proven by antidepressant response predictions in recent clinical tests (Wu et al., 2020).

Another innovative point in the study is dlPFC function. Recently, more and more studies have identified the key role of dlPFC in top-down cognitive-emotional control of the affective state, generating structures as well as process of attention responses (Notzon, Steinberg, Zwanzger, & Junghofer, 2018). For a long time, the dlPFC has been assumed to realize its executive function for affective states and subsequent behavior via dorsal system. For instance, pioneer studies showed hypoactivation of the right dlPFC during negative material processing in major depressive disorder (MDD) (Zhong, Pu, & Yao, 2016). In addition, the dlPFC send signals to its adjacent regions such as the orbitofrontal cortex, dorsomedial prefrontal cortex, and hippocampus, to elicit an appropriate response (Ghashghaei & Barbas, 2002; Kringelbach, 2005; Rangel & Hare, 2010). In situations where mood and emotion are involved, the dlPFC is always activated, especially during tasks that require direct attention. Notably, these findings have advanced the neurobiological understanding of the dlPFC’s function during emotional attention.

As mentioned previously, psychiatric disorders with aberrant emotional processes are likely to show some symptoms that are characteristic of lesions in the dlPFC (Phillips, Drevets, Rauch, & Lane, 2003). For instance, previous studies indicated reduced cortical neuronal size and reduced glial cell density in schizophrenic patients, as well as reduced glial cell number and prefrontal cortex volumes in bipolar disorder patients (Ketter et al., 2001). Furthermore, studies in MDD patients pointed at the bilateral dlPFC hypoactivations during episode, mainly reflect in reduced density and dlPFC volumes along with reductions in metabolism and blood flow employing functional neuroimaging techniques (Rajkowska et al., 1999).

Except for these structural and functional abnormalities, dlPFC impairment is also reflected as abnormal neural oscillations, especially the alpha response. Recent research has explored alpha activity in the dlPFC activity (i.e., repetitive transcranial magnetic stimulation [rTMS], transcranial direct current stimulation [tDCS], and tACS) (Ironside et al., 2019).

Recent rTMS research demonstrated that excitatory rTMS induced in the dlPFC strengthened top-down control of aversive facial stimuli and induced effects on emotional arousal of fearful facial in MDD patients (Notzon et al., 2018). Thus, low-field magnetic stimulation synchronized to the IAF is thought to be more effective for MDD treatment (Leuchter et al., 2015). Additionally, anodal tDCS in the left dlPFC has resulted in obvious effects on emotion regulation. Another research study in our lab has discovered the emotional effects on alpha asymmetry, which has not yet been published. Thus far, few studies have tested the emotional or attentional effects of alpha-tACS in the dlPFC.

Although participants did not report any behavioral or psychological anomalies during the experiment, we cannot entirely exclude the differences in mood or other emotional characteristics between two groups, which would influence the results. However, most of the participants were university students, which could have weakened the influence of these factors.

Considering these limitations, the results demonstrated the entrainment of individual alpha-tACS and its essential role in emotional attention, at least in healthy subjects. However, it is not clear to what extent tACS can influence this function. Systematic comparison between other neurostimulation techniques for exact mechanism as well the combination of functional imaging techniques, which offer excellent resolution in both temporal and spatial domains, are still lacking.

The presented findings offer important insights into the crucial roles of prefrontal alpha measures and dlPFC function in the area of emotional attention. The revealing of such effects should encourage more research in order to explore the intrinsic mechanisms influencing the dlPFC and tACS as well their neurophysiologic correlates. With additional research, the neurophysiological profiles of emotion processing may serve as valuable biomarkers for diagnoses and reveal potential target brain regions for therapeutic interventions in various neuropsychiatric disorders.

## Acknowledgments

This work was funded by supported by National Natural Science Foundation of China (No.81630051 and No.81801786). No parts of the manuscript and data have been published previously. The authors would like to thank all of the subjects for their participation in the study.

## Author Contributions

Shuang Liu and Yuchen He contributed equally to this work and should both be considered as first authors.

## Appendix

## Description of active and sham tACS settings

In the active group, total active stimulation required 20 minutes. In the sham group, only a short period of tACS was administered with 10 cycles ramp-up and 10 cycles ramp-down. This ensure that participants in the sham group experienced a slight acupuncture feeling, similar to the active group. However, participants in the sham group continued wearing the electrode cap after the stimulation phase to experience stimulation for the same duration. Stimulation frequency and intensity setting were the same in both groups.

## EEG preprocessing

Preprocessing was performed offline using EEGLAB in the MATLAB (R2013b) environment, which is an open-source MATLAB-based toolbox for data processing of electrophysiological signals. The reference electrode was first adjusted to the binaural mean (M1 and M2) to avoid differences between the left and right hemispheres. Then, the raw resting EEG signals were down-sampled to 500 Hz and bandpass-filtered from 0.5 to 120 Hz, while down-sampled the task EEG signals to 200 Hz and bandpass-filtered from 0.5 to 45 Hz with a 50 Hz notch filter for both, and followed by independent component analysis (ICA) to remove artifacts caused by heartbeats and eye movements.

## IAF detection

The resting EEG of “eyes open” durations were segmented into 1 second epochs. FFT spectrums were calculated for each epoch separately, then averaged per epoch. The prominent alpha peak of the averaged spectrum (8–13 Hz) at electrode F3/F4 was detected as IAF, ranging from 8–11 Hz across participants.

## ERP Acquisition

Raw EEG data was divided into three emotional states. To safely acquire complete ERP waveforms, the epoch for ERP was 950 ms from 250 ms pre-stimulus to 700 ms post-stimulus. Data was then subjected to 1–30 Hz bandpass filtering to observe the waveform. All of the epochs were baseline-corrected using data from the first 200 ms pre-stimulus. Trials with amplitudes greater than 100μV were discarded by subtracting the minimum amplitude by maximum amplitude. Ultimately, grand averaging of dataset responses to three types of emotional stimuli resulted in three ERP waveforms for each participant (target, distracter, and neutral) within the cranial channels of interest.

